# The genetic basis of hindwing eyespot number variation in *Bicyclus anynana* butterflies

**DOI:** 10.1101/504506

**Authors:** Angel G. Rivera-Colón, Erica L. Westerman, Steven M. Van Belleghem, Antónia Monteiro, Riccardo Papa

## Abstract

The underlying genetic changes that regulate the appearance and disappearance of repeated traits, or serial homologs, remain poorly understood. One hypothesis is that variation in genomic regions flanking master regulatory genes, also known as input-output genes, controls variation in trait number, making the locus of evolution almost predictable. Other hypotheses implicate genetic variation in up-stream or downstream loci of master control genes. Here, we use the butterfly *Bicyclus anynana*, a species which exhibits natural variation in eyespot number on the dorsal hindwing, to test these two hypotheses. We first estimated the heritability of dorsal hindwing eyespot number by breeding multiple butterfly families differing in eyespot number, and regressing eyespot number of offspring on mid-parent values. We then estimated the number and identity of independent genetic loci contributing to eyespot number variation by performing a genome-wide association study with restriction site-associated DNA Sequencing (RAD-seq) from multiple individuals varying in number of eyespots sampled across a freely breeding lab population. We found that dorsal hindwing eyespot number has a moderately high heritability of approximately 0.50. In addition, multiple loci near previously identified genes involved in eyespot development display high association with dorsal hindwing eyespot number, suggesting that homolog number variation is likely determined by regulatory changes at multiple loci that build the trait and not by variation at single master regulators or input-output genes.

## Introduction

Body plans often evolve through changes in the number of repeated parts or serial homologs by either addition or subtraction. For instance, the pelvic fins of vertebrates are inferred to have originated after the appearance of pectoral fins, perhaps via co-option of the pectoral or caudal fin developmental programs to a novel location in the body (Larouche, Zelditch, & Cloutier, 2017; Ruvinsky & Gibson-Brown, 2000). In insects, the absence of limbs and wings in the abdomen is inferred to be due to the repression (or modification) of limbs and wings in these segments by *hox* genes (Galant & Carroll, 2002; Ohde, Yaginuma, & Niimi, 2013; Ronshaugen, McGinnis, & McGinnis, 2002; Tomoyasu, Wheeler, & Denell, 2005). Regulatory targets of abdominal *hox* genes are likely to underlie loss of limb/wing number in these body segments (Ohde et al., 2013; Tomoyasu et al., 2005), although these mutations have not yet been identified. Thus, while serial homolog number variation is a common feature in the evolution of organisms’ body plans, the underlying genetic changes that regulate the appearance and disappearance of these repeated traits remain poorly understood.

Studies in *Drosophila* have contributed most to the identification of the genetic basis underlying the evolution of serial homolog number. Larvae of different species have different numbers of small hairs, or trichomes, in their bodies, and variation in regulatory DNA around the gene *shavenbaby* appears to be largely responsible for this variation (McGregor et al., 2007). Moreover, *shavenbaby* has been labeled a master regulatory gene because its ectopic expression in bare regions of the body leads to trichomes (Payre, Vincent, & Carreno, 1999). However, a more complex genetic architecture seems to underlie variation in the number of larger bristles found in the thorax of adults. In this case variation around *achaete-scute*, a gene complex required for bristle differentiation, plays a role in controlling bristle number variation across species (Marcellini & Simpson, 2006). Interestingly, genetic variation in upstream regulatory factors, whose spatial expression overlaps some, but not all bristles, is also known to impact bristle number in lab mutants (Garcia-Bellido & de Celis, 2009). Finally, *shavenbaby* and *scute* genes are also known as input-output genes due to their central “middle of the hour-glass” position in regulatory networks (Stern & Orgogozo, 2008). These genes respond to the input of multiple upstream protein signals, present at distinct locations in the body, and in turn control the regulation of the same battery of downstream genes, to affect the same output (trichome or bristle development) at each of these body locations. Mutations in the regulatory regions of these genes are thus expected to have minimal pleiotropic effects, and to lead to changes in the number of times the network is deployed, and thus to evolution in the number of trichome or bristles in the bodies of these flies. While this type of regulatory network architecture points to predictable regions in the genome that will readily evolve leading to trait number evolution, i.e., hotspots of evolution, it might represent only one type of architecture among others that are still unexplored. More systems, thus, need to be investigated for a more thorough understanding of the genetic basis underlying variation of repeated traits in bodies.

One promising system for investigating the genetic basis of serial homolog number evolution are the eyespot patterns on the wings of nymphalid butterflies. Eyespots originally appeared on the ventral hindwing in a lineage of nymphalid butterflies, sister to the Danainae, and have subsequently been added to the forewings and dorsal surfaces of both wings (Oliver, Beaulieu, Gall, Piel, & Monteiro, 2014; Oliver, Tong, Gall, Piel, & Monteiro, 2012; Schachat, Oliver, & Monteiro, 2015). Furthermore, within a single species, eyespot number can vary significantly between individuals or sexes (Brakefield & van Noordwijk, 1985; Owen, 1993; Tokita, Oliver, & Monteiro, 2013), allowing for population genetic approaches to identify the underlying genetic basis of such variation. Genes controlling eyespot number variation within a species might also be involved in promoting this type of variation seen across species.

One of the best model species for studying the genetic basis of eyespot number variation is the nymphalid butterfly *Bicyclus anynana*. This species exhibits natural variation and sexual dimorphism in eyespot number on the dorsal hindwing surface, which play a possible role in mate choice (Westerman, Chirathivat, Schyling, & Monteiro, 2014). The observed variation consists of males averaging 0.75 dorsal hindwing eyespots, with a range of 0-3, and females averaging 1.5 dorsal hindwing eyespots, with a range of 0-5 (Westerman et al., 2014) (Fig. 1). Lab populations of this species also display a series of mutant variants that affect eyespot number on other wing surfaces. Genetic and developmental studies on eyespot number variation in this species suggest the existence of at least two different underlying molecular mechanisms. Spontaneous mutants such as Spotty (Brakefield & French, 1993;

**Figure 1.**
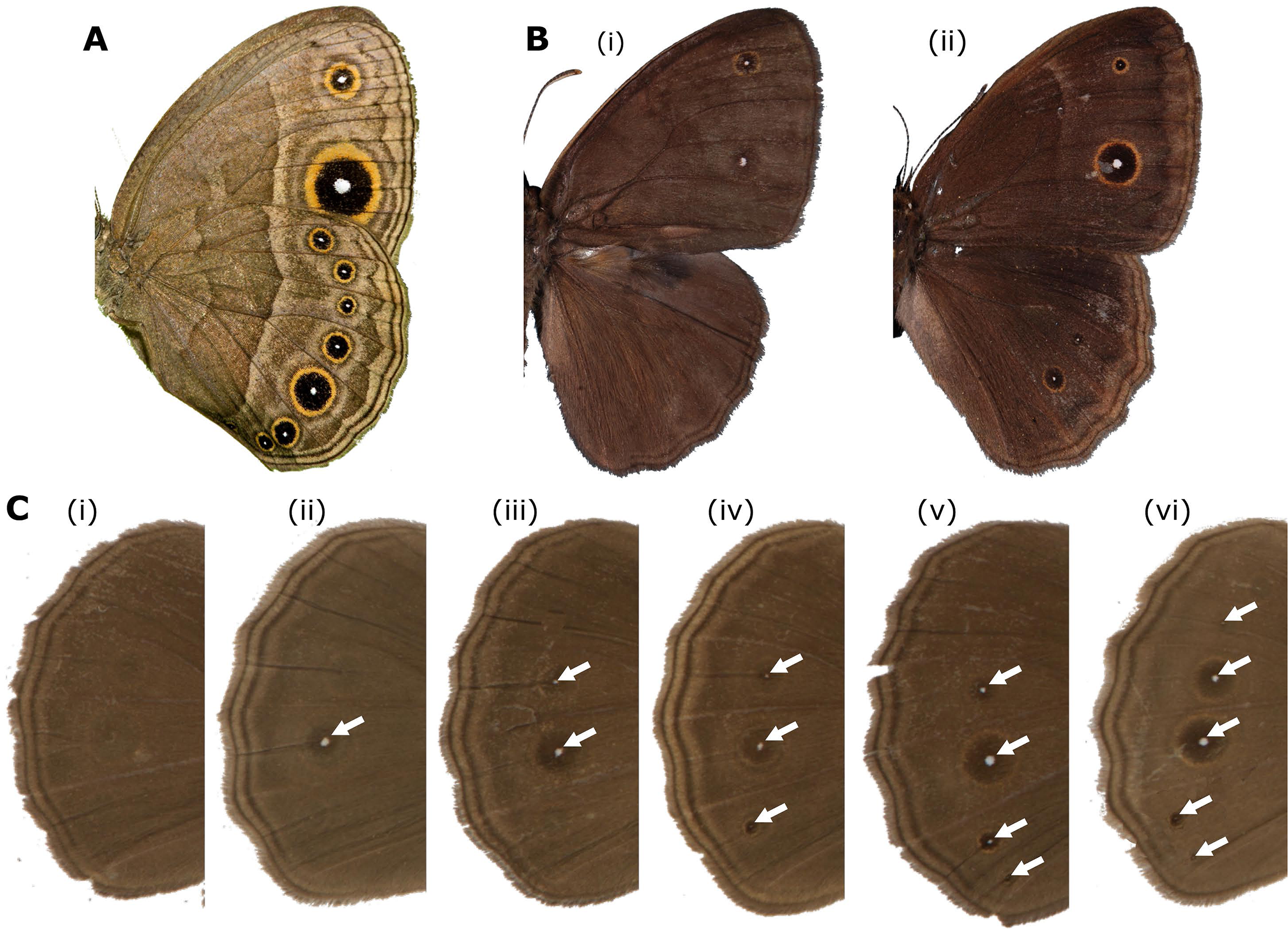
Eyespot pattern and number variation in *Bicyclus anynana.* (A) Eyespot pattern on the ventral side of wings: forewing displays two eyespots; hindwing displays seven eyespots. (B) Eyespots pattern on dorsal side of wings. Male (left) displaying two dorsal forewing eyespots and zero dorsal hindwing eyespots and female displaying two dorsal forewing eyespots and three dorsal hindwing eyespots. (C) Dorsal hindwing eyespot number (DHEN) variation, ranging from zero to five UV-reflective spots, marked by white arrows (i–vi).

Monteiro et al., 2013; Monteiro, Brakefield, & French, 1997), Missing (Monteiro et al., 2007), P- and A- (Beldade, French, & Brakefield, 2008), or X-ray induced mutations such as 3+4 (Monteiro, Prijs, Bax, Hakkaart, & Brakefield, 2003), segregate as single Mendelian alleles, and cause discrete and obvious changes in eyespot number or affect the size of very specific eyespots. On the other hand, multiple alleles of small effect likely regulate the presence or absence of small eyespots that sometimes appear between the typical two eyespots on the forewing, or on the most posterior wing sector of the ventral hindwing. This type of eyespot number variation is positively correlated with eyespot size variation, responds readily to artificial selection on eyespot size (Beldade & Brakefield, 2003; Holloway, Brakefield, & Kofman, 1993; Monteiro, Brakefield, & French, 1994), and is likely under the regulation of a threshold type mechanism (Brakefield & van Noordwijk, 1985).

Interestingly, eyespot number variation within *B. anynana* can involve changes to single eyespots or to several eyespots at a time, on one or both wing surfaces. For instance, Spotty introduces two eyespots on the dorsal and ventral surfaces of the forewing, whereas A- and P-primarily reduce the size of the single anterior (A-) or the posterior eyespot (P-) of the dorsal surface exclusively, without affecting eyespot size or number on the ventral surface. The genetic basis for these differences is still unknown.

Recently, the gene *apterousA* (*apA*) was shown to regulate wing pattern differences between dorsal and ventral surfaces in *B. anynana*, including differences in eyespot number (Prakash & Monteiro, 2018). This gene is expressed exclusively on the dorsal wing surfaces and its mutation via CRISPR-Cas9 led to dorsal wing surfaces acquiring a ventral identity, which included additional eyespots. This study indicated that *apA* is a repressor of eyespots on the dorsal surface. However, *B. anynana*, has eyespots on dorsal wing surfaces and their presence and variation in number appears to be correlated with variation in the number of small circular patches, positioned at future eyespot centers, lacking *apA* expression (Prakash & Monteiro, 2018). This suggests that genetic variation at loci that modulate the expression of *apA* in eyespot centers on the dorsal surface, or genetic variation in regulatory regions of *apA* itself, might be involved in regulating eyespot number specifically on the dorsal surface of wings.

The genetic architecture of eyespot number variation in any butterfly species remains unknown. Here, we examine the genetic basis of dorsal hindwing eyespot number (DHEN) variation in *B. anynana*. We carried out two sets of experiments. We first estimated the heritability for this trait by breeding multiple butterfly families differing in eyespot number and regressing eyespot number of offspring on mid-parent values. Then we estimated the number and identity of independent genetic loci that are contributing to variation in this trait by performing a genome-wide association study with restriction site-associated DNA Sequencing (RAD-seq) from multiple individuals varying in number of eyespots sampled across a freely breeding lab population.

## Materials and Methods

### Study organism

*Bicyclus anynana* is a Nymphalid butterfly common to sub-tropical Africa for which a colony has been maintained in the laboratory since 1988. All *Bicyclus anynana* butterflies used in this study were collected from a colony established in New Haven, CT (Yale University), composed of an admixed population of numerous generations of freely breeding individuals with variable dorsal hindwing eyespot number phenotypes. Individuals from this colony originated from an artificial colony established in Leiden University in 1988, which was established from numerous gravid females collected in Malawi in 1988. Previous studies have estimated that this laboratory population maintains genetic diversity comparable to those of natural populations (reviewed in Westerman et al., 2016). The colony was kept in controlled conditions of 12 hours light/dark cycles, 80% relative humidity and a temperature of 27^°^C. Larvae were fed on corn plants and adult butterflies on mashed banana, as described in previous publications (Westerman et al., 2014).

### Heritability of dorsal hindwing eyespot number

We examined the number of dorsal hindwing eyespots (DHEN) on all offspring from 18 separately reared families whose parents differed in eyespot number: six families where both parents had DHEN of zero (0F x 0M); six where both parents had DHEN of one (1F x 1M); and six where both parents had DHEN of two (2F x 2M). All generations were reared in the conditions described above. We ensured virginity of the females by separating the butterflies in the parental generation into sex-specific cages on the day of eclosion. All families were started within 5 days of each other using adults ranging from 1-3 days old (ANOVA, n=18, DF=2, F=0.8266, p=0.4565). Each breeding pair was placed in a cylindrical hanging net cage of 30 cm diameter X 40 cm height, with food (banana slices), water and a young corn plant on which to lay eggs. When corn plants were covered with eggs, they were placed in family-specific mesh sleeve cages for larval growth. Females were given new plants on which to lay eggs until they died. Pupae and pre-pupae were removed from the sleeve cages and placed in family-specific cylindrical hanging net cages for eclosion. The cages were checked daily for newly emerged butterflies. On the day of eclosion, DHEN was recorded for each offspring. Heritability was calculated by regressing offspring on midparent values, correcting the estimate for assortative mating, as described in Falconer & Mackay (1996). Estimates were obtained for the pooled offspring data as well as for separate regressions of female and male offspring data on mid-parent values. Sex-specific heritabilities were calculated using the correction for unequal variances in the two sexes (Falconer and Mackay, 1996). We then tested for an interaction of parental phenotype and offspring sex on offspring phenotype using a general linear model, with sex, parental phenotype, and sex*parental phenotype as fixed variables.

### Sample collection and phenotype determination for genomic association study

To identify regions in the genome that are associated with DHEN variation, we collected and sequenced a total of 30 individuals. Fifteen individuals contained no eyespots (absence) and 15 containing two or more eyespots (presence) (Table S1). Both groups contained an assorted number of male and female individuals. Wings of the collected individuals were removed, and the bodies were preserved in ethanol for DNA extraction.

### RAD library preparation and sequencing

Genomic DNA of the preserved bodies was extracted using DNeasy Blood & Tissue Kit (Qiagen), with an additional RNase digestion for removing RNA from the extracted nucleic acid samples. The quality and concentration of the extracted DNA was verified using gel electrophoresis and Qubit 2.0 fluorometer (Life Technologies). Extracted genomic DNA was used for preparing Illumina RAD sequencing libraries based on previously described protocols (Baird et al., 2008; Etter, Bassham, Hohenlohe, Johnson, & Cresko, 2011). DNA was digested with the frequent cutting enzyme *Pst*1 and ligated to P1 adapters containing a unique barcode 5 bp in length. Samples were pooled and sheared using a Covaris M220 (Covaris Inc.) instrument and size selected for 300-500 bp inserts on average. After end-repair and P2 adapter ligation, the library was amplified by PCR. The pooled library was then sequenced utilizing a single lane of an Illumina HiSeq2000® 100 bp paired-end module.

### Read quality and filtering

Following sequencing of a RAD-seq library composed of 30 *B. anynana* individuals, we obtained 127 million paired-end reads 100 bp in length. The raw RAD reads were demultiplexed using *Stacks v1.42* (Catchen, Hohenlohe, Bassham, Amores, & Cresko, 2013; Catchen et al., 2011) *process_radtags* pipeline, and reads with low quality and/or ambiguous barcodes were discarded. Further, we removed Illumina adapter sequences from the reads and trimmed sequences to 80 bp in length, as suggested from the FastQC quality control tool (http://www.bioinformatics.babraham.ac.uk/projects/fastqc/) results. We retained a total of 111 million (86 %) and an individual average of 3.7 million ± 1.2 million filtered paired-end reads.

### Reference alignment

The 111 million retained read pairs were aligned to the most recent *B. anynana* genome assembly (v1.2) with its corresponding annotation (Nowell et al., 2017). This reference assembly is composed of 10,800 individual scaffolds, for a total genome size of about 475 Mb (N50 = 638.3 kb). The annotation contains 22,642 genes, with a partial CEGMA completeness of 97.2 %. Filtered reads were aligned to the reference genome using BWA v0.7.13 (H. Li & Durbin, 2009) *mem* with default seed lengths, mismatch and gap scores, but allowing for the marking of shorter split reads as secondary alignments for compatibility with PicardTools v1.123 (https://broadinstitute.github.io/picard/). Resulting alignments were directly converted to BAM files using SamTools v1.12 (H. Li et al., 2009) *view*. BAM files were then sorted with SamTools *index* and filtered for duplicates using PicardTools v1.123 *MarkDuplicates* and processed with *AddOrReplaceReadGroups* for GATK compatibility. In total, we obtained a 92.9 % read alignment and 79.6 % properly mapped read pairs. At each RAD locus, average per-individual sequencing coverage was 18.8X (± 4.3; median: 18.3).

### Variant calling and association mapping

Identification of associated regions of the genome was performed simultaneously using two different analytical approaches to produce method-agnostic results. The first one, referred to as the “association method”, identified variants on the filtered BAM files using GATK v3.5 (McKenna et al., 2010) *UnifiedGenotyper* with the default call confidence values and outputting only variant sites. These variant sites were then filtered using VCFTools v0.1.14 (Danecek et al., 2011) *recode* to obtain only calls with a minimum genotype quality of 30, a minimum genotype depth of 5, present in over 50 % of all individuals, a max allele number of 2, and minimum allele frequency of 0.05. This results in a filtered VCF containing only high-quality biallelic variants. After genotyping with *GATK*, we obtained 350,121 filtered SNPs. Genotype-to-phenotype association in the genotyped samples via the “association method” was performed using PLINK v1.90 (Clarke et al., 2011; Purcell et al., 2007). We used a genome-wide adjusted Fisher analysis to identify genotype-to-phenotype association values for each SNP. The test also implements an adaptive Monte Carlo permutation analysis to reduce the detection of false positives. This process tests the obtained *p*-values per locus after each successive permutation, with a maximum of 1,000,000 replications. The second analytical approach, referred to as the “*F_ST_* method”, called SNPs using Stacks v1.42. Using the *ref_map.pl* wrapper script with default parameters, RAD loci were assembled from the reference genome-mapped reads using *pstacks*. A catalog of all loci was generated with *cstacks* and samples were matched to this catalog using *sstacks.* The *populations* program was then run on this catalog to generate population genetic measures, enabling the calculation of F-statistics. For a variant to be included in the analysis, it had to be present in both study groups in over 75% percent of individuals (p=2, r=0.75) and have a minimum allele frequency above 0.05. Additionally, a Fisher’s Exact Test p-value correction was applied to the resulting *F_ST_* and Analysis of Molecular Variance (AMOVA) *F_ST_* values as a multiple testing correction. Using *Stacks* for the F_ST_ method we reconstructed 207,752 RAD loci and 673,340 raw SNPs. After filtering, 73,159 RAD loci and 238,786 SNPs were retained. Filtered variants obtained from both datasets were compared to retain only SNPs with support from both genotyping methods. A total of 216,338 SNPs were shared between the two datasets and were used for subsequent comparisons.

### Selection of candidate loci

To minimize the identification of false genotype-to-phenotype relationships, only areas of the genome displaying both association and *F_ST_* peaks were used for further analysis. The use of both metrics simultaneously also ensured that the relationships observed are method agnostic. This multi-metric approach, including a combination of association and *F_ST_* outliers, has been utilized repeatedly to identify genomic regions associated with domestication in both dogs and cats (Axelsson et al., 2013; Montague et al., 2014), variation in feather coloration in warblers (Brelsford, Toews, & Irwin, 2017), and the architecture and modularity of wing pattern variation in *Heliconius* butterflies (Nadeau et al., 2014; Van Belleghem et al., 2017). Significant peaks were defined as areas of the genome with SNPs containing association and/or *F_ST_* 10 standard deviations above the genome-wide mean. Candidate variants were then annotated using *SnpEff* v4.3T (Cingolani et al., 2012), building a *de novo* database with the available *B. anynana* reference annotation, to identify possible effects over nearby genes.

### Principal Component Analysis

To determine the baseline-level genome-wide diversity and divergence among individuals in the present population, we performed a Principal Component Analysis (PCA) on the obtained genotypes. Although our sampled individuals originated from a single, freely breeding population, performing this analysis allowed us to corroborate that the observed genomic diversity lacks any substructuring that could impact our outlier identification. To do this, we randomly selected a subset of 5000 filtered variants from the *Stacks* catalog and made them into a whitelist, as described by the *Stacks* manual and by Rochette & Catchen (2017). We then ran the *populations* module on this subset of variants with the addition of the *--genepop* export format flag. The resulting genepop file was processed using the *adegenet* v2.1.1 R package (Jombart, 2008; Jombart & Ahmed, 2011) by converting the genotype calls into a *genind* object, scaling missing data by mean allele frequency and analyzed with PCA.

### Ordering of the *B. anynana* scaffolds along the *Heliconius melpomene* genome

The current *B. anynana* reference assembly (Nowell et al., 2017) has an N50 of 638.3 kb and is composed of 10,800 unlinked scaffolds. To assess whether associated SNPs on separate *B. anynana* genome scaffolds could be part of the same block of association, we ordered the scaffolds of the *B. anynana* genome along the *Heliconius melpomene* v2 genome assembly (Davey et al., 2016). Although *B. anynana* and *H. melpomene* diverged about 80 My ago (Espeland et al., 2018) and have a different karyotype (n=28 in *B. anynana* versus n=21 in *H. melpomene*), the *H. melpomene* genome is the most closely related butterfly genome that has been assembled into highly contiguous chromosomal scaffolds using pedigree informed linkage maps. Aligning both genomes provides valuable information to interpret our association analysis. To construct this alignment, we used the alignment tool *promer* from the MUMmer v3.0 software suite (Kurtz et al., 2004). *Promer* was used with default settings to search for matches between sequences translated in all six possible reading frames between the *B. anynana* and *H. melpomene* genome. The obtained alignments were subsequently filtered for a minimum alignment length of 200 bp and a minimum percent identity (%IDY = (matches x 100)/(length of aligned region)) of 90 %. These filtered alignments were used to order the *B. anynana* scaffolds according to the order in which they aligned along the *H. melpomene* genome. If a scaffold aligned to multiple locations or chromosomes, priority was given to the position it matched with highest identity. For scaffolds that contained significant associations with hindwing eyespot number, we also retained alignments with a minimum %IDY of 70 % and a minimum alignment length of 150 bp to investigate possible fine scale rearrangements between the *B. anynana* and *H. melpomene* genome.

### Linkage disequilibrium analysis

In addition to ordering the *B. anynana* scaffolds to the *H. melpomene* genome for assessing the genomic linkage of SNPs, we calculated linkage disequilibrium in our *B. anynana* study population. To calculate linkage disequilibrium for genomic SNPs, we phased 213,000 SNPs that were genotyped in all samples using *beagle* v4.1 (Browning & Browning, 2007). Estimates of linkage disequilibrium were calculated from 100,000 randomly selected SNPs, using the *VCFtools* v0.1.14 (Danecek et al., 2011) *– hap-r2* function, with a max LD window of 5 Mbp, and minimum allele frequency cutoff of 0.10. Resulting LD comparisons for genomic SNPs were then plotted in R, where a Loess local regression was calculated and used to determine the genome-wide window size of linkage disequilibrium decay. Subsequently, this LD window size was used for the investigation of genes near associated loci.

## Results

### Dorsal hindwing spot number variation has moderate to high heritability

Zero spot females were only produced by 0×0 families and one 1×1 family, and were absent from any 2×2 DHS families. Zero spot males, however, were produced by all 0×0 families, all but one of the 1×1 families and all but one of the 2×2 families. Two spot females were produced by all but one (a 0×0) family, while 2 spot males were produced by all 2×2 families, but only two 1×1 families and one 0×0 family (Table 1). These results demonstrate that alleles are sufficiently segregating in our experimental design to perform heritability estimates.

**Table 1.** DHEN is heritable. Summary DHEN data for offspring from 6 families each of 0×0, 1×1, and 2×2 DHEN crosses, separated by sex. Offspring with asymmetric DHEN are included xg11in the average DHEN estimate.

|  | **0 x 0 DHEN Families** |  | **1 x 1 DHEN Families** |  | **2 x 2 DHEN Families** |  |
| --- | --- | --- | --- | --- | --- | --- |
|  | Females | Males | Females | Males | Females | Males |
| 0 DHEN | 46 | 159 | 2 | 47 | 0 | 14 |
| 1 DHEN | 55 | 15 | 45 | 36 | 19 | 38 |
| 2 DHEN | 24 | 2 | 43 | 9 | 67 | 27 |
| 3 DHEN | 2 | 0 | 4 | 0 | 6 | 0 |
| **Average DHEN** | **0.87** | **0.17** | **1.49** | **0.63** | **1.87** | **1.13** |

DHEN has a heritability of 0.4442 ± 0.264 for females, 0.5684 ± 0.306 for males, and 0.5029 ± 0.159 when the sexes were pooled. There was no significant interaction of parental phenotype and offspring sex on offspring phenotype (General linear Model with sex, parental phenotype, and sex*parental phenotype as parameters, AICc (Akaike Information Criterion) = 1498.204, effect tests: sex χ^2^=293.361, p<0.0001; parental phenotype χ^2^=271.56, p<0.0001; sex* parental phenotype χ^2^=0.032, p=0.8576).

### Genome-wide variation and linkage disequilibrium of the study population

To confirm the absence of population substructure in our study population, we calculated measures of genetic variation and diversity between the samples displaying presence of DHEN (pre) and samples with absent DHEN (abs). As expected, the two groups showed very little genome-wide genetic divergence, with a genome-wide *F_ST_* equal to 0.0075. The absence of any population substructure between the two sampled phenotype groups was further demonstrated by complete overlap of the two groups in the PCA as well as little contribution of phenotype group to the observed variation in first and second Principal Components (Fig. 2A). Additionally, we observed very similar genome-wide nucleotide diversity in the group displaying presence of DHEN (pre,π = 0.0090) when compared to the group with absent DHEN (abs, π = 0.0083). Hence, we do not observe any demographic substructuring of the study population that could potentially bias our genetic association analysis.

**Figure 2.**
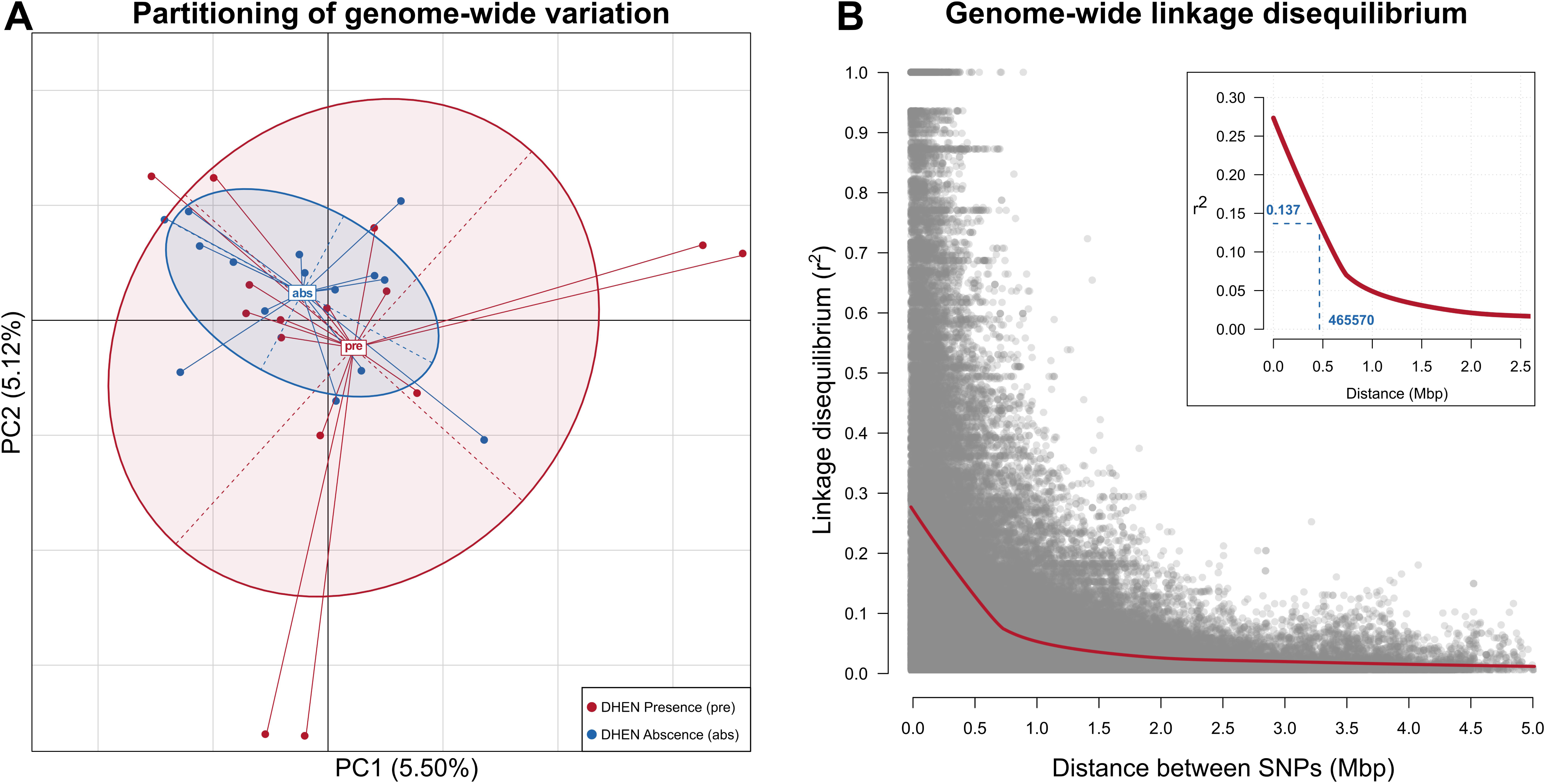
Genome-wide population structure and linkage disequilibrium (LD) in the study population. **(A.)** Principal Component Analysis (PCA) of the allelic variation observed in 5000 randomly-selected genome-wide SNPs in the study population across both phenotype groups, DHE presence (*pre*, red) and DHE absence (*abs,* blue). Ellipses display boundaries of the 95% confidence interval. Little contribution to variation in the principal components and overlap of the variation on both phenotype groups suggests lack of underlying demographic substructuring in the study population. **(B.)** Genome-wide linkage disequilibrium (LD) in the *B. anynana* study population. Grey dots represent LD values for a SNP pairwise comparison. In red, Loess regression smoothed curve representing LD decay. Insert: Zoomed-in LD decay curve, indicating distance at which LD is halved (465,570 bp) and corresponding r^2^ value (0.137).

After calculating genome-wide estimates of linkage disequilibrium decay (Fig. 2B), we observed a max smoothed r^2^ value of 0.272, and a halving of r^2^ within 464 Kb (Fig. 2B). This window size suggests that average linkage blocks are around 500 Kb in length, and that variants within this distance are in strong linkage disequilibrium.

### Association mapping of dorsal hindwing spot number variation

After mapping and genotyping RAD loci across the *B. anynana* reference (Nowell et al., 2017), we identified a total of 216,338 SNPs shared between the two different genotyping strategies used (see methods), of which 340 SNPs display both elevated *F_ST_* and significantly high association with DHEN variation. These candidate SNPs are located in 15 different scaffolds of the *B. anynana* genome assembly spanning 3.54 Mbp equal to 0.744% of the whole genome (Fig. 3). The relatively close proximity of associated variants within the 15 associated scaffolds, particularly within our 500 kb LD windows, suggests that only 15 discrete regions influence DHEN variation. Further ordering these scaffolds along the contiguously assembled *Heliconius melpomene* genome suggest that they likely belong to 10 to 11 different genomic regions in the genome (Fig. 4) and thus strongly suggest DHEN variation is a polygenic trait with multiple loci.

**Figure 3.**
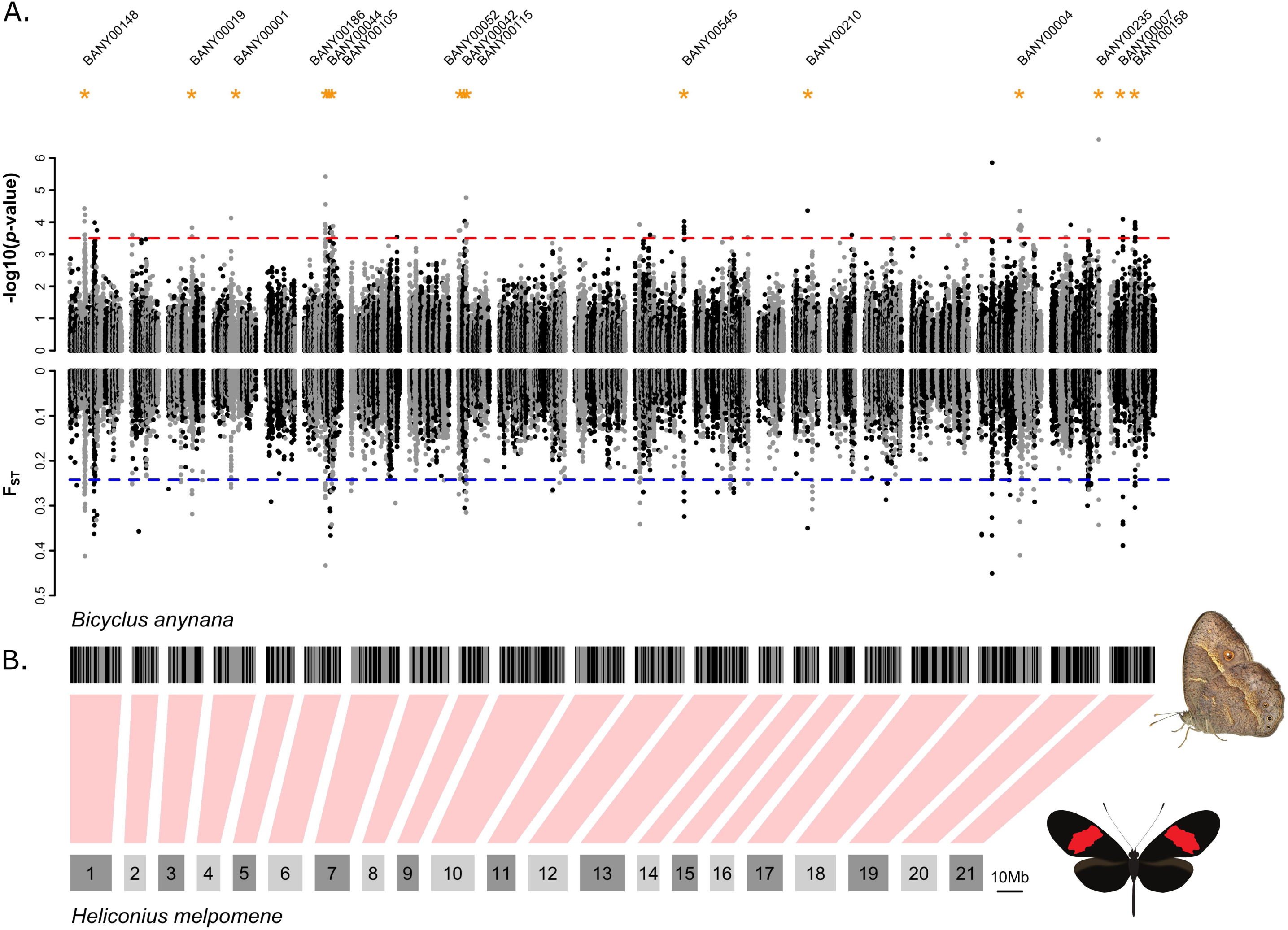
Genome-wide association with dorsal hindwing eyespot number. **(A.)** Plots show genomic association to dorsal hindwing eyespot number (top) and *F_ST_* between individuals with different dorsal hindwing eyespot numbers (bottom). Each dot represents a single SNP. Dashed lines represent the threshold for detecting a significant genome-wide association (top, in red) and *F_ST_* (bottom, in blue). Scaffolds containing both significant association and *F_ST_* outliers are marked with asterisks. **(B.)** Genomic scaffolds from the *Bicyclus_anynana*_v1x2 genome are arranged along the 21 chromosomes of the *Heliconius melpomene* v2 assembly. For ordering the *B. anynana* scaffolds along the *H. melpomene* genome, only matches with a minimum percentage of identity of 90% and a minimum alignment length of 200 bp were used. If scaffolds matched multiple *H. melpomene* chromosomes, the scaffold was positioned along the chromosome to which it had the most matches. Using this strategy 76.7% of the *B. anynana* genome scaffolds were aligned to the *H. melpomene* genome.

**Figure 4.**
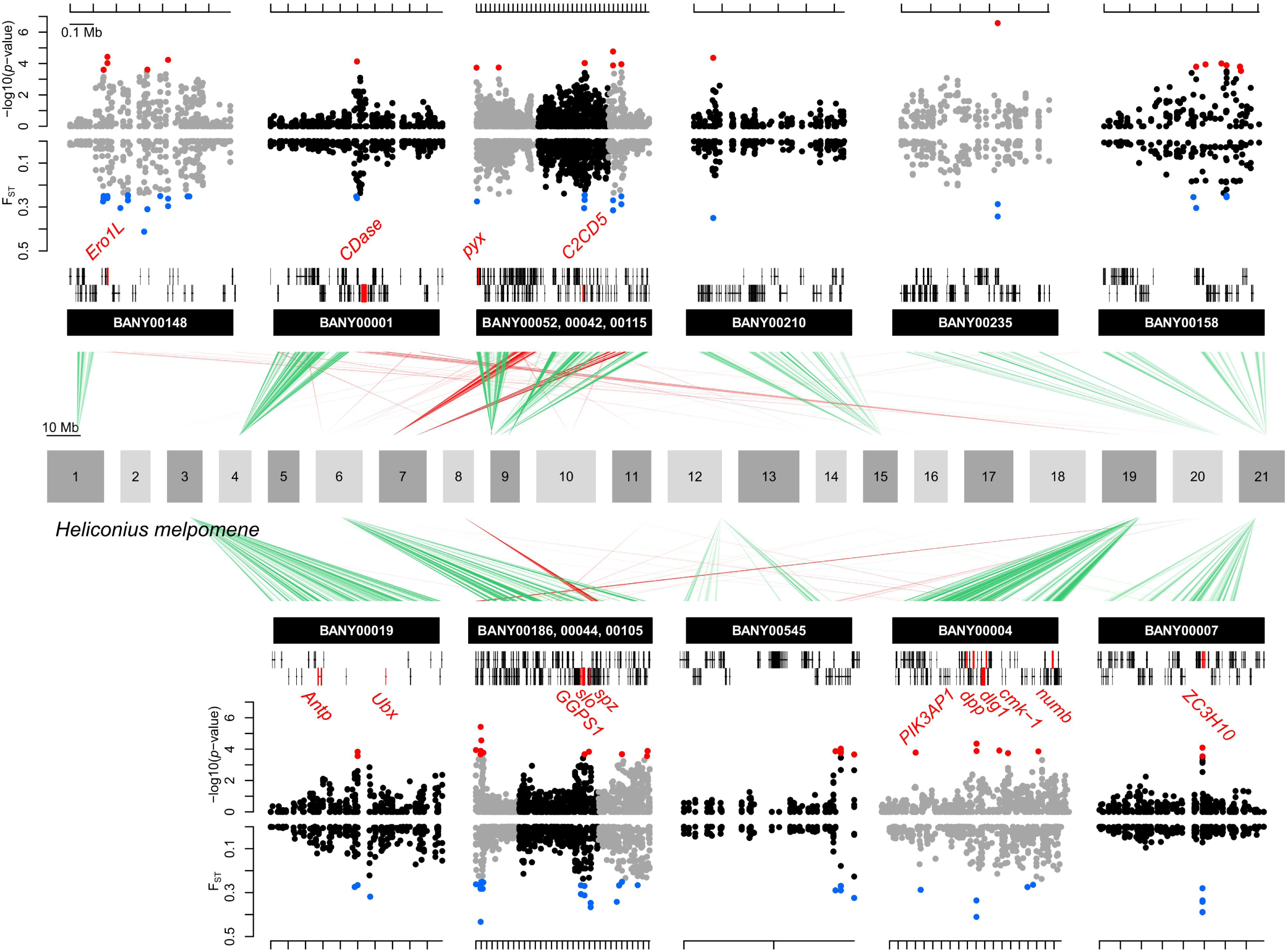
Zoom-in on putative genomic regions underlying dorsal hindwing eyespot number variation. Plots show genomic association to dorsal hindwing eyespot number (top) and *F_ST_* (bottom) between individuals with different dorsal hindwing eyespot numbers for scaffolds with significant outliers (red for association and blue for *F_ST_*). Each dot represents a single SNP. Green and red lines show matches of the *B. anynana* scaffolds (minimum percentage of identity of 70% and a minimum alignment length of 150 bp) to the *H. melpomene* v2 assembly. Green lines represent the most frequent matches of the scaffold to a *H. melpomene* chromosome, whereas red lines represent matches to a different *H. melpomene* chromosome. Vertical black rectangles represent gene models. Gene models in red represent genes that have previously been demonstrated to be involved in eye spot development.

### Candidate gene identification

Using the available Lepbase reference annotation for the *B. anynana* v1.2 assembly we identified the neighboring annotated genes and relative positioning of the 340 outlier SNPs common to both the *F_ST_* and genome-wide association. The majority of these SNPs were in non-coding sequence, with 116 (34.1%) being intergenic, 78 (22.9%) immediately upstream, and 49 (14.4%) immediately downstream of annotated genes (Supplemental Table S3). Of the 340 outlier SNPs, 11 (3.2%) produce non-synonymous changes to coding regions, while 1 (0.3%) causes change in a splice region sequence.

When the annotation is observed at a genic level, the biggest proportion of SNPs occur closely downstream or upstream of annotated genes, 10 (23.8%) and 12 (28.6%) genes, respectively, and could have regulatory effects over nearby genes (Supplemental Table S3). The 11 SNPs that cause non-synonymous changes are located within only two genes: 1) BANY.1.2.g01110 (Zinc finger CCCH domain-containing protein 10, *ZC3H10*) has a Serine to Cysteine substitution in position 114, and a Isoleucine to Valine substitution in position 142. 2) BANY.1.2.g04901 (Geranylgeranyl pyrophosphate synthase, *GGPS1*) contains a Arginine to Glutamine substitution, and a Isoleucine to Threonine substitution in positions 96 and 98, respectively. Finally, BANY.1.2.g13875 (Acidic fibroblast growth factor intracellular-binding protein, *Fibp*), contains the identified splice region variant.

A number of annotated genes observed within or nearby associated regions of the genome were implicated in eyespot development in previous studies, whereas other genes are here implicated for the first time (Table 2). BANY.1.2.g00030 (Neutral Ceramidase, *CDase*) was previously identified as differentially expressed in eyespots relative to flanking wing tissue (Özsu & Monteiro, 2017). BANY.1.2.g04910 (Protein spaetzle, *spz*) is involved in the regulation of the Toll signaling pathway, recently implicated in eyespot development (Özsu & Monteiro, 2017). Calcium signaling-related genes were also identified, including BANY.1.2.g00659 (Calcium/calmodulin-dependent protein kinase type 1, *cmk-*1), BANY.1.2.g04904 (Calcium-activated potassium channel slowpoke, *slo*) and BANY.1.2.g05412 (Transient receptor potential channel pyrexia, *pyx*). Calcium signaling has been recently associated with eyespot formation in *B. anynana* (Özsu & Monteiro, 2017). BANY.1.2.g10819 (Ero1-like protein, *Ero1L*), BANY.1.2.g00658 (Disks large 1 tumor suppressor protein, *dlg1*) and BANY.1.2.g00681 (Protein numb, *numb*) have functions related to Notch signaling, a pathway previously associated with eyespot formation (Reed & Serfas, 2004). Additionally, BANY.1.2.g02571 (Homeotic protein antennapedia, *Antp*) known to be expressed in early stages of eyespot development (Saenko, Marialva, & Beldade, 2011) and BANY.1.2.g00653 (Protein decapentaplegic, *dpp*) expressed in dynamic patterns during eyespot center formation during the larval stage (Connahs et al., 2017 - *bioRxiv*; Monteiro, Glaser, Stockslager, Glansdorp, & Ramos, 2006) were also identified within areas of high association. BANY.1.2.g04715 (C2 domain-containing protein 5, *C2CD5*) appears to be involved in insulin receptor signaling, a pathway that has, so far, not been implicated in eyespot development nor eyespot plasticity.

**Table 2.** DHEN candidate genes. BANY.1.2 gene ID, gene name, molecular function, biological process, BANY.1.2 scaffold ID all refer to the *B. anynana* v1.2 genome assembly and annotation (Nowesll et al., 2017).

| **BANY.1.2 gene ID** | **Gene name** | **Gene description** | **BANY.1.2 scaffold ID** | **Known candidate** |
| --- | --- | --- | --- | --- |
| g00030 | *CDase* | Neutral ceramidase | BANY00001 | Yes (Özsu & Monteiro, 2017) |
| g00647 | *PIK3AP1* | Phosphoinositide 3-kinase adapter protein 1 | BANY00004 | No |
| g00653 | *dpp* | Protein decapentaplegic | BANY00004 | Yes (Connahs et al., 2017 - *bioRxiv*) |
| g00658 | *dlg1* | Disks large 1 tumor suppressor protein | BANY00004 | Yes (Cheah, Chia, & Yang, 2000; Q. Li et al., 2009; Tien et al., 2008) |
| g00659 | *cmk-1* | Calcium/calmodulin-dependent protein kinase type 1 | BANY00004 | No |
| g00681 | *numb* | Protein numb | BANY00004 | Yes (Cheah, Chia, & Yang, 2000; Q. Li et al., 2009; Tien et al., 2008) |
| g01110 | *ZC3H10* | Zinc finger CCCH domain-containing protein 10 | BANY00007 | No |
| g02571 | *Antp* | Homeotic protein antenapedia | BANY00019 | Yes (Saenko, Marialva, & Beldade, 2011) |
| g02579 | *Ubx* | Homeotic protein ultrabithorax | BANY00019 | Yes (Weatherbee 1999; Tomoyasu et al. 2005; Tong et al. 2014) |
| g04715 | *C2CD5* | C2 domain-containing protein 5 | BANY00042 | No |
| g04901 | *GGPS1* | Geranylgeranyl pyrophosphate synthase | BANY00044 | No |
| g04904 | *slo* | Calcium-activated potassium channel slowpoke | BANY00044 | No |
| g04910 | *spz* | Protein spaetzle | BANY00044 | No |
| g05412 | *pyx* | Transient receptor potential channel pyrexia | BANY00052 | No |
| g10819 | *Ero1L* | Ero1-like protein | BANY00148 | Yes (Cheah, Chia, & Yang, 2000; Q. Li et al., 2009; Tien et al., 2008) |

## Discussion

The use of population genomic analyses, including association mapping, GWAS, and *F_ST_* scans, has been extensively used in natural populations to identify the genomic components underlying a number of biological processes such as hybridization and speciation, local adaptation, and ecological and landscape genomics (Reviewed in Campbell, Poelstra, & Yoder, 2018; Narum, Buerkle, Davey, Miller, & Hohenlohe, 2013; Rellstab, Gugerli, Eckert, Hancock, & Holderegger, 2015). In our work, we applied a population genomics approach to identifying the genetic component of dorsal hindwing number variation in the butterfly *B. anynana* via the comparison of genomic diversity between individuals from a single laboratory-maintained, free-breeding population. We demonstrated that free-breeding in the laboratory population has likely maintained homogenization of the genetic variation across the genome due to a lack of demographic substructuring, reducing the detection of false positive associations between SNPs and the trait of interest, DHEN variation.

The combined results of our heritability and genome-wide association study suggest that variation in dorsal hindwing eyespot number in *B. anynana* is a complex trait regulated by multiple loci. Using a combination of RAD sequencing and genome-wide association mapping, we identified 10 to 11 potentially distinct genomic regions in the *B. anynana* reference genome associated with variation in DHEN. Further analysis of these genomic regions highlights a total of 15 candidate genes (Table 2). Some (7) of these genes have previously been found associated with eyespot development via their expression patterns or via functional studies, while others (8) are suggested for the first time. Complete list of all annotated genes identified in the associated regions is provided in the supplemental data (Supplemental Table S2).

### DHEN variation and known elements of the eyespot regulatory network

Among our identified candidates, we observe a number of genes previously implicated in eyespot development in *B. anynana*. Our analysis suggests six of the genes known to be expressed in eyespots are associated with DHEN: *Ubx, Antp, Dpp, dlg1, numb*, *Ero1L*, and *CDase*. BANY.1.2.g02579 is annotated as the hox protein Ultrabithorax (*Ubx*), an important selector gene that gives insect hindwings, including those of butterflies, a different identity from forewings (Tong, Hrycaj, Podlaha, Popadić, & Monteiro, 2014; Weatherbee et al., 1999). In *B. anynana,* this gene is expressed across the whole hindwing, a conserved expression pattern observed across insects, but has additional, stronger, expression in the eyespot centers of the hindwing only, something that is not seen in other butterflies with eyespot such as *Junonia coenia* (Tong et al., 2014). Over-expression of this gene led to eyespot size reductions in both fore and hindwings of *B. anynana* (Tong et al., 2014). However, absence of *Ubx* in clones of cells in the hindwing of *J. coenia* (via a spontaneous unknown mutation) (Weatherbee et al., 1999) and CRISPR knockouts in *B. anynana* (Y. Matsuoka, unpublished) led to both eyespot enlargements (of Cu1 eyespots to sizes that match Cu1 forewing eyespot sizes), and to complete deletions of eyespots that are normally present in the ventral hindwing (M2 and M3) but absent on the forewing. These results suggest both a repressing role and an activating role for *Ubx* that depends on eyespot position on the hindwing. Our results indicate that genetic variation at *Ubx* might contribute to eyespot number variation either via a threshold-like mechanism acting on eyespot size or a more discrete mechanism regulating presence or absence of eyespots in specific wing sectors.

BANY.1.2.g02571 is annotated as the protein Antennapedia (*Antp*), another hox gene involved in the differentiation of the anterior-posterior body axis in insects and involved in the differentiation of the thoracic limbs in *Bombyx* moths (Chen et al., 2013). *Antp* expression has been observed as one of the earliest expressed genes in developing eyespot centers (Saenko et al., 2011), and in dorsal eyespots (Özsu & Monteiro, 2017). Knock-out of *Antp* indicates this gene is required for forewing eyespot development and for the development of the white centers and the full size of the hindwing eyespots (Y. Matsuoka, unpublished). Variation at this locus appears to contribute to hindwing eyespot number variation, perhaps using a similar threshold-like mechanisms to that proposed for *Ubx*, based on eyespot size regulation.

BANY.1.2.g00653 is annotated as decapentaplegic (*Dpp*), a candidate morphogen likely involved in the differentiation of the eyespot centers in *B. anynana* via a reaction-diffusion mechanism (Connahs et al., 2017 - *bioRxiv*). *Dpp* mRNA is expressed in regions around the developing eyespots in mid to late larval wings (Connahs et al., 2017 - *bioRxiv*) in anti-colocalized patterns to Armadillo, a transcription factor effector of the Wingless signaling pathway, expressed in the actual eyespot centers in late larval stages (Connahs et al., 2017 - *bioRxiv*). Genetic variation either linked with the protein coding sequence of *Dpp*, or more likely with its regulatory region affects eyespot number variation in hindwings.

Three genes known to interact with another eyespot-associated gene, *Notch* (Reed & Serfas, 2004), are also implicated in our study. BANY.1.2.g00658 (Disks large 1 tumor suppressor protein, *dlg1*), BANY.1.2.g00681 (protein numb, *numb*), and BANY.1.2.g10819 (Ero1-like protein, *Ero1L*) are known to interact and regulate the *Notch* (*N*) signaling pathway in *Drosophila melanogaster* (Cheah, Chia, & Yang, 2000; Q. Li et al., 2009; Tien et al., 2008). The existence and role of these interactions are unknown in *B. anynana*, as is the role of the *Notch* receptor itself. However, the *Notch* receptor has a dynamic pattern of expression (Reed & Serfas, 2004) that is very similar to that of *Distal-less*, a gene that has recently been implicated in setting up the eyespot centers likely via a reaction-diffusion mechanism (Connahs et al., 2017 - *bioRxiv*). Genetic variation at these three genes could be interacting with the eyespot differentiation process through unknown mechanisms.

Newly identified components of pathways previously associated with eyespot development, such as Toll and Calcium signaling (Özsu & Monteiro, 2017), have also been observed among our candidates. BANY.1.2.g00959 (Calcium/calmodulin-dependent protein kinase type 1, *cmk-1*), BANY.1.2.g04715 (C2 domain-containing protein 5, *C2CD5*), BANY.1.2.g04904 (Calcium-activated potassium channel slowpoke, *slo*) and BANY.1.2.g05412 (Transient receptor potential channel pyrexia, *pyx*) all possess Calcium binding and/or interactions with Calcium signaling among their annotated functions, while BANY.1.2.g00647 (Phosphoinositide 3-kinase adapter protein 1, *PIK3AP1*) and BANY.1.2.g04910 (Protein spaetzle, *spz*) both interact and/or regulate the Toll signaling pathway among their annotated functions. *Spaetzle*, in particular, is a ligand that enables the activation of the Toll pathway in *Drosophila* (Yamamoto-Hino & Goto, 2016). The role of *Spaetzle* is currently unknown in the context of eyespot development but this ligand could be an interesting target of further study. Our data suggests that genetic variation at these loci are also regulating hindwing eyespot number variation.

### Effects of non-coding mutations in the evolution of eyespot number variation

After identifying 10 to 11 regions of the *B. anynana* genome associated with DHEN variation and characterizing the relationship of identified SNPs with nearby genes we observe that the majority of the SNPs fall outside coding sequences (90.8%), and most of the ones that do fall inside protein coding sequences result in synonymous mutations (Supplemental figure/table). Only two genes, BANY.1.2.g01110 (Zinc finger CCCH domain-containing protein 10, *ZC3H10*) and BANY.1.2.g04901 (Geranylgeranyl pyrophosphate synthase, *GGPS1*) contain SNPs that represent non-synonymous mutations of unknown effect. Such non-coding DNA variation linked to DHEN is likely to be *cis*-regulatory and controlling the expression of the nearby genes described above. *cis-*regulatory elements are thought to have profound implications in the evolution of morphological diversity (Carroll, 2008). Particularly, they have been associated with variation in pigmentation patterns in a wide variety of animal systems, including the evolution of eggspot pigmentation patterns in cichlids (Santos et al., 2014), wing pigmentation patterns in *Drosophila* (Koshikawa et al., 2015; Werner, Koshikawa, Williams, & Carroll, 2010), divergent pigmentation patterns in capuchino seedeater finches (Campagna et al., 2017), variation in red, black and yellow color patterns in *Heliconius* butterflies due to regulatory changes in the *optix, WntA* and *Cortex* genes (Martin & Reed, 2014; Reed et al., 2011; Supple et al., 2013; Van Belleghem et al., 2017) among others. In the case of eyespot number variation, regulatory mutations around the genes identified here might disrupt the reaction-diffusion mechanism of eyespot center differentiation (Connahs et al., 2017 - *bioRxiv*), or later processes of eyespot center signaling, that eventually translate to presence of absence of an eyespot in particular wing sectors.

Concluding, the genetic variation uncovered in this work affects eyespot number variation on the dorsal surface but not on the ventral surface of the wing. Thus, our work suggests that the genetic variants identified with our analysis affect eyespot number in a surface-specific manner. This surface-specific regulation is potentially mediated via *apA*, a previously identified dorsal eyespot repressor (Prakash & Monteiro, 2018). The polygenic nature of our results argue that genetic variation at the loci identified above, e.g., *Antp, Ubx*, *dpp*, etc, rather than at the *apA* locus itself regulates dorsal eyespot number. The way in which these genes interact is unclear, but changes in gene expression at the identified loci might impact the repression of *apA* in specific wing sectors on the dorsal surface, allowing eyespots to differentiate in those sectors.

Finally, the use of a genome-wide sequencing strategy allowed us to discover a series of independent loci that appear to contribute to DHEN in *B. anynana*. These loci, predominantly composed of polymorphisms in non-coding DNA, suggest that changes in DHEN are mostly occurring in regions that regulate the expression of previously known eyespot-associated genes. Thus, while our work has enriched the list of genes involved in eyespot number variation, it also confirms that variation at multiple genes, rather than at a single top master regulator or input-output gene (such as *shavenbaby* or *achaete-scute*) is involved in regulating number of serial homologs. This highlights a more complex, but still poorly understood, genetic architecture for serial homolog number regulation.

## Acknowledgements

We thank Elizabeth Schyling for help rearing families for heritability analysis, and Robert Rak and Chris Bolick for rearing corn plants for the larvae. We also thank the UPR-RP Sequencing and Genomics Facility and the UPR-RP High Performance Computing Facility for additional support in library preparation and computational analysis. This work was funded by a NSF IOS-110382 Doctoral Dissertation Improvement Grant to ELW and AM, a NSF PR-LSAMP Bridge to the Doctorate Program (NSF Grant Award HRD1139888) to AGRC, and a Ministry of Education, Singapore grant (MOE2015-T2-2-159) to AM.

